# Genome-wide association study and low-cost genomic predictions for growth and fillet yield in Nile tilapia (*Oreochromis niloticus*)

**DOI:** 10.1101/573022

**Authors:** Grazyella M. Yoshida, Jean P. Lhorente, Katharina Correa, Jose Soto, Diego Salas, José M. Yáñez

## Abstract

Fillet yield (FY) and harvest weight (HW) are economically important traits in Nile tilapia production. Genetic improvement of these traits, especially for FY, are lacking, due to the absence of efficient methods to measure the traits without sacrificing fish and the use of information from relatives to selection. However, genomic information could be used by genomic selection to improve traits that are difficult to measure directly in selection candidates, as in the case of FY. The objectives of this study were: (i) to perform genome-wide association studies (GWAS) to dissect the genetic architecture of FY and HW, (ii) to evaluate the accuracy of genotype imputation and (iii) to assess the accuracy of genomic selection using true and imputed low-density (LD) single nucleotide polymorphism (SNP) panels to determine a cost-effective strategy for practical implementation of genomic information in tilapia breeding programs. The data set consisted of 5,866 phenotyped animals and 1,238 genotyped animals (108 parents and 1,130 offspring) using a 50K SNP panel. The GWAS were performed using all genotyped and phenotyped animals. The genotyped imputation was performed from LD panels (LD0.5K, LD1K and LD3K) to high-density panel (HD), using information from parents and 20% of offspring in the reference set and the remaining 80% in the validation set. In addition, we tested the accuracy of genomic selection using true and imputed genotypes comparing the accuracy obtained from pedigree-based best linear unbiased prediction (PBLUP) and genomic predictions. The results from GWAS supports evidence of the polygenic nature of FY and HW. The accuracy of imputation ranged from 0.90 to 0.98 for LD0.5K and LD3K, respectively. The accuracy of genomic prediction outperformed the estimated breeding value from PBLUP. The use of imputation for genomic selection resulted in an increased relative accuracy independent of the trait and LD panel analyzed. The present results suggest that genotype imputation could be a cost-effective strategy for genomic selection in tilapia breeding programs.

## INTRODUCTION

Investment in selective breeding programs generates economic return because genetic selection is aimed at improving productivity of important economic traits that result in permanent, cumulative and sustainable changes in a farm’s population over generations of selection (Gjedrem 2012). In a simulation study, Ponzoni et al. (2007) estimated that the benefit/cost ratio reached a maximum of 60/1 with the implementation of family based breeding programs in Nile tilapia. Therefore, selective breeding is an important tool to increase aquaculture production and profitability, satisfying the increasing demand for animal protein (Gjedrem 2012).

The first Nile tilapia breeding program was established in 1988 and since then high levels of genetic gains have been achieved for economically important traits, e.g. genetic gains for body weight ranged from 7 to 20% per generation (Bentsen et al., 2017; Eknath et al., 1998; Gjedrem et al., 2012 and Khaw et al., 2008). However, until now the Nile tilapia breeding programs have been based only on pedigree and phenotype information for genetic evaluations. The incorporation of genomic information for genetic analysis has not been evaluated or implemented in tilapia breeding programs. This is mainly due to the fact that dense SNP panels were not available until recently (Joshi *et al.* 2018; Yáñez *et al.* 2019). The use of genomic information for the implementation of genomic selection has already been assessed in various aquaculture species, e.g. Atlantic salmon, rainbow trout, salmon coho, common carp, channel catfish and Pacific oyster (Barria et al., 2018a; Garcia et al., 2018; Gutierrez et al., 2018; Palaiokostas et al., 2018; Vallejo et al., 2018; Yoshida et al., 2018a; Bangera et al., 2017; Correa et al., 2017; Meuwissen et al., 2014; Ødegård et al., 2014; Tsai et al., 2016). As it has been demonstrated in these studies, an increase in selection accuracy when including genomic information from dense SNP panels, especially for traits which are difficult to measure in selection candidates (Yañez and Martinez 2010; Yáñez *et al.* 2014). Carcass quality traits (e.g. fillet yield) are considered key traits in the breeding goal for Nile tilapia genetic improvement (Nguyen *et al.* 2010; Ponzoni *et al.* 2011) and these traits could be more efficiently improved through the inclusion of genomic information in genetic evaluations.

The use of genomic information from dense SNP panels provides the opportunity to increase the rate of genetic progress in breeding programs (Meuwissen *et al.* 2001). However, the cost of genotyping is high and alternative methods are necessary for cost-efficient genomic applications (VanRaden, *et al.* 2011; Carvalheiro *et al.* 2014). Strategies such as selective genotyping (Sen *et al.* 2009; Jiménez-Montero *et al.* 2012; Ødegård and Meuwissen 2014), genotyping animals using low-density panels (Tsai *et al.* 2016; Bangera *et al.* 2017; Correa *et al.* 2017; Yoshida *et al.* 2018a) and genotype imputation (Cleveland and Hickey 2014; Sargolzaei *et al.* 2014; Chen *et al.* 2014) have been tested as alternative strategies for reducing costs for the practical implementation of genomic information in breeding programs.

Imputation of genotypes reduces the cost of genomic selection, by genotyping a small proportion of animals (e.g. parents or influential animals) using a dense SNP panel and selection candidates using a LD SNP panel, and then imputing (predicting) missing genotypes from the lower to the HD SNP panel (Sargolzaei *et al.* 2009). In aquaculture species, these cost-effective strategies have been assessed and reported to generate genomic prediction accuracies similar to those obtained when all selection candidates are genotyped with HD SNP panels (Dufflocq et al., 2019; Tsai et al., 2017; Yoshida et al., 2018b).

The objectives of this study were: (i) to perform a genome-wide association study to dissect the genetic architecture and identify molecular markers for growth and fillet yield; (ii) to evaluate the accuracy of genotype imputation as an alternative low-cost genotyping strategy, and (iii) to assess the accuracy of genomic selection for growth and fillet yield using true and imputed SNP genotypes in farmed Nile tilapia. To our knowledge, this is the first study evaluating the incorporation of true and imputed dense genotypes for the implementation of low-cost genomic predictions in farmed Nile tilapia.

## MATERIAL AND METHODS

### Phenotypes

The Nile tilapia population used in the current study belongs to a breeding nucleus established by Aquacorporación Internacional group (GACI) in Costa Rica. The origin of the population is described in detail by Yoshida et al. (2019). Briefly, the eggs of each full-sib family were incubated and reared in separate hapas until tagging. A mating design of two dams per sire was used to produce full and half-families, which varied from 74 to 89 families across the year-classes (Table 1). For each year-class an average number of 18 fish/family (ranging from 5 to 49) were tagged and reared until they were an average of 13 months old, where the traits fillet yield (FY) and harvest weight (HW) were recorded for each individual fish.

**Table 1.** Summary statistics for phenotyped animals by year-class.

| Year-class | Fam<sup>1</sup> | Genotyped | Age |  | Fillet Yield |  |  | Harvest Weight |  |  |
| --- | --- | --- | --- | --- | --- | --- | --- | --- | --- | --- |
|  |  |  | Mean | SD<sup>2</sup> | Number | Mean (%) | SD<sup>2</sup> | Number | Mean (g) | SD<sup>2</sup> |
| **2011** | 89 | - | 376.25 | 24.24 | 1,004 | 36.34 | 1.85 | 1,027 | 919.15 | 257.61 |
| **2012** | 82 | - | 343.48 | 16.33 | 0760 | 34.47 | 2.05 | 0767 | 730.79 | 235.62 |
| **2013** | 80 | - | 514.22 | 14.49 | 2,628 | 34.07 | 2.45 | 2,636 | 907.91 | 268.04 |
| **2017** | 74 | 1,130* | 370.54 | 20.04 | 1,474 | 31.74 | 2.16 | 1,479 | 953.57 | 252.86 |

\*The parents of year-class 2017 were genotyped using 50K SNPs panel to perform genotype imputation.
<sup>1</sup>Family number;
<sup>2</sup>Standard deviation.

### Genotypes

Genomic DNA was extracted from fin clip samples from 108 parents (45 sires and 63 dams) and 1,364 offspring from year-class 2017. Samples were then genotyped using a 50K SNP Illumina BeadChip, which is described in detail by Yáñez et al. (2019). Genomic DNA was purified from all the samples using the DNeasy Blood & Tissue Kit (QIAGEN) according to the manufacturer’s protocol. Before the genome-wide association study (GWAS) and imputation analysis, genotypes and samples were filtered according to the following exclusion criteria: Hardy-Weinberg Disequilibrium (HWE, p-value < 1×10^-6^), Minor Allele Frequency (MAF < 0.05) and genotyping call-rate for SNP and samples < 0.95.

### Genome-wide association analysis

We performed the GWAS to dissect the genetic architecture and to identify regions of the Nile tilapia genome containing SNPs with important effects on FY and HW. We used the weighted single step genomic best linear unbiased prediction (wssGBLUP) method (Wang *et al.* 2012) implemented in postGSf90 module from BLUPf90 family programs (Misztal *et al*. 2016). The following model was used:

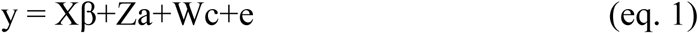

where *y* is a vector of phenotypes (FY or HW), β is a vector of contemporary group as fixed effects that comprise the year-class:sex:tank, and harvest weight or age for FY and HW as covariate, respectively; a is a vector of random additive direct genetic effects; c is a vector of common environmental effect and e is a vector of residual effect. X, Z and W are incidence matrices for β, a and c effects, respectively.

The wssGBLUP is similar to the pedigree-based BLUP (PBLUP) method except for the use of a combined genomic and pedigree relationship. The kinship matrix A^-1^ is replaced by matrix H^-1^ (Aguilar *et al.* 2010), which combines genotype and pedigree relationship coefficients:

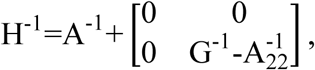

where, 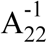 is the inverse of a pedigree-based relationship matrix for genotyped animals; and G^-1^ is the inverse genomic relationship matrix. The SNPs were assumed with an initial value of one corresponding to the single-step genomic BLUP (ssGBLUP) method (Misztal *et al.* 2009). In the wssGBLUP the marker variances were estimated from allele frequencies and used as weights, which were updated on each iteration (Wang *et al.* 2014). We tested three iterations of weights, and used the second iteration to plot the manhattans plots, as suggested by Wang et al. (2012) and Zhang et al. (2016), who proposed that two iterations were sufficient to correctly identify major SNPs in wssGBLUP. The wssGBLUP included all HD genotyped animals (parents and offspring) which passed quality control (n = 1,238) and all the phenotyped fish present in Table 1.

To evaluate the presence of putative genes associated with the traits under study, we reported all genes between the first and last SNP position of each 20-SNP window, searched using BLAST (*Basic Local Alignment Search Tool*) of the SNP probes against the last version of the *Oreochromis niloticus* reference genome (Conte *et al.* 2018), publicly available at NCBI (GenBank assembly accession GCA_001858045.3).

### Genotype imputation

Three *in silico* LD panels were constructed with SNP densities of 500 (LD0.5K), 1,000 (LD1K) and 3,000 (LD3K). The SNPs from the LD panels were selected using the option --indep-pairwise of Plink v1.9 software (Purcell *et al.* 2007), with a window size of 180,432 kb, a step of 1 SNPs and a variable r^2^ according to chromosome. This command produced a subset of markers that are proportional for chromosome size, are approximately evenly spaced and in low linkage disequilibrium with each other as recommended by Cleveland and Hickey (2014).

The imputations were run using genotype information from 108 parents and 20% of the offspring (n = 225) as a reference set (HD panel) and 80% of offspring (n = 904) were used as the validation set (LD panel). The assignment of the offspring to the reference and validation sets was random, and five replicates were used each time. In addition, we used pedigree information available for all individuals for imputation, consisting of eight generations of records for each genotyped animal. Imputation of genotypes was performed using the FImpute v2.2 software (Sargolzaei et al., 2014) and the accuracy of imputation was calculated as the correlation between true and imputed genotypes for the validation set.

### Genomic selection

We used true and imputed genotypes from the three LD panels (LD0.5K, LD1K and LD3K) to assess prediction accuracies using a five-fold cross validation scheme. In addition, accuracy of breeding values was also estimated using pedigree-based information (PBLUP method) and the true 32K SNPs (true markers that passed in quality control). Briefly, all genotyped animals (n = 904) with phenotypes were randomly divided into five exclusive training sets (80% of the dataset; n = 721 and SD = 5 animals) which were used to estimate the SNP effects; the remaining animals were used as validation sets (20% of the dataset; mean = 193 and SD = 5 animals), for which their phenotypes were masked and their performance was predicted based on the marker effects. This five-fold cross validation was replicated five times for each SNP panel density and the results are presented as a mean for all replications.

We used the BLUPF90 family of programs (Misztal *et al.* 2016) to perform the genetic evaluations using pedigree-based information and the ssGBLUP method which uses both pedigree and genomic information, and additional information of the animals with only phenotypes (Table 1) in the validation set. The statistical model fitted was the same of the eq. 1, except for PBLUP method, for which the kinship matrix used was A^-1^ instead of H^-1^ in ssGBLUP.

Prediction accuracies were calculated in the validation sets using the following equation:

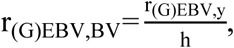

where r_GEBV,y_ is the correlation between the estimated breeding value (EBV) or genomic estimated breeding values (GEBV) of a given model (predicted for the validation set using information from the training set) and the true phenotypic record, while *h* is the square root of the pedigree-based estimate of heritability.

### Genetic parameters and heritability

The total additive genetic variance 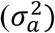 was estimated using the kinship matrix A and H for PBLUP and ssGBLUP, respectively. For all traits studied, the heritabilities were computed using the following equation:

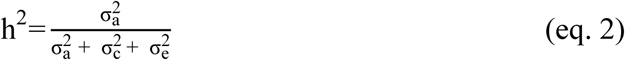

where, 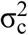 and 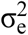 is the common environmental and residual variance, respectively.

### Data availability

All raw phenotypic and genotypic data used in the current study can be found at the Figshare public repository (https://figshare.com/s/9b265a22b7e138c5a839).

## RESULTS

### Basic statistics and genotype quality control

The total number of individuals phenotyped ranged from 5,866 to 5,909 for FY and HW, respectively, and varied per year-class with the maximum number of animals phenotyped in 2013. On average, the recorded fish were 401 days old at harvest weight. The average FY was 34.2% (SD = 2.13% g) and the average HW was 878 g (SD = 254 g) for phenotyped fish (Table 1).

Out of the initial 1,364 individuals and 43,272 SNPs which were effectively genotyped, a total of 1,130 animals and 32,306 SNPs (32K) passed quality control. The MAF < 0.05 parameter excluded the highest number of SNPs (∼ 4.8K), whereas HWE and genotyped call-rate excluded about 2K and 4K SNPs, respectively.

### Genetic parameters and heritability

For both FY and HW the additive genetic variance and heritability were slightly higher when using genomic information compared to the pedigree-based method. For instance, heritability values using ssGBLUP were 0.21 and 0.36 for FY and HW, respectively. For PBLUP heritability for FY and HW was estimated to be 0.21 to 0.31, respectively. Additionally, a reduction in error of heritability estimates was shown for ssGBLUP when compared with PBLUP.

### Genome-wide association analysis

Manhattan plots for the proportion of genetic variance explained by each 20-SNP window for FY and HW are shown in Figures 1 and 2. A total of 1,624 20-SNP windows with average length of 530 kb (range from 10 to 6,690 kb) were obtained. After the second iteration of wssGBLUP, the top five windows cumulatively explained 5.2 and 8.0% of the total genetic variance for FY and HW, respectively (Table 2).

**Table 2.**
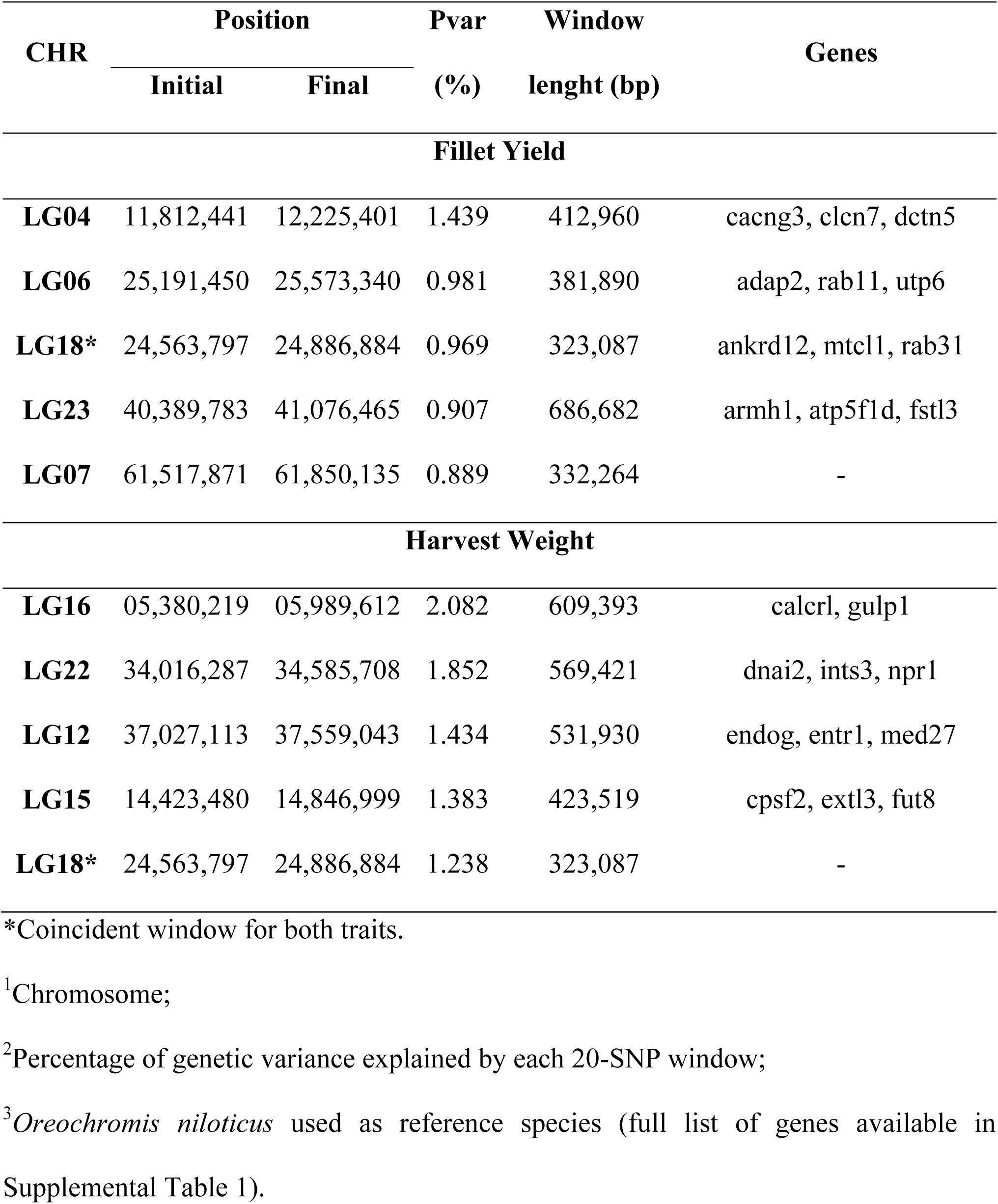
Top five ranked 20-SNP windows that explain the largest proportion of genetic variance for fillet yield and harvest weight in Nile tilapia.

**Table 3.** Estimates of variance components and heritability for fillet yield and harvest weight in Nile tilapia.

| Traits | PBLUP |  |  |  |  | ssGBLUP* |  |  |  |  |
| --- | --- | --- | --- | --- | --- | --- | --- | --- | --- | --- |
| | $\sigma_a^{21}$ | $\sigma_c^{22}$ | $\sigma_e^{23}$ | $h^2^4$ | $SE^5$ | $\sigma_a^{21}$ | $\sigma_c^{22}$ | $\sigma_e^{23}$ | $h^2^4$ | $SE^5$ |
| **Fillet yield** | 0.972 | 0.174 | 3.498 | 0.209 | 0.053 | 0.972 | 0.168 | 3.491 | 0.210 | 0.038 |
| **Harvest weight** | 19,312 | 4,296 | 39,522 | 0.306 | 0.073 | 23,161 | 3,823 | 37,345 | 0.360 | 0.047 |

\*Estimated for true 32K genotype panel.
<sup>1</sup>Additive genetic variance;
<sup>2</sup>Common environment variance;
<sup>3</sup>Residual variance;
<sup>4</sup>Heritability;
<sup>5</sup>Standard error.

**Figure 1.**
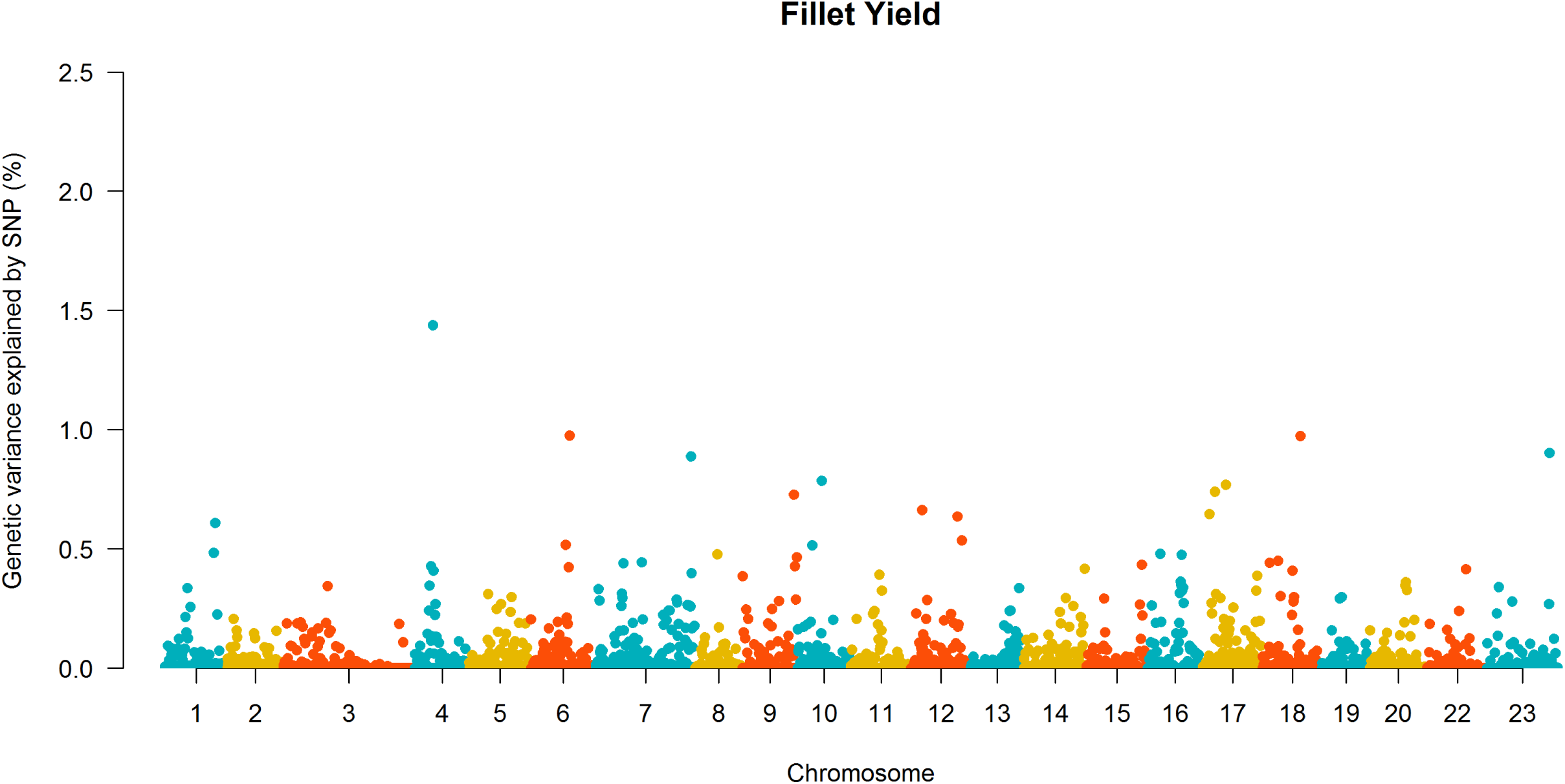
Manhattan plot of genetic variance explained by 20-SNP windows for fillet yield in the 2^nd^ iteration of wssGBLUP.

**Figure 2.**
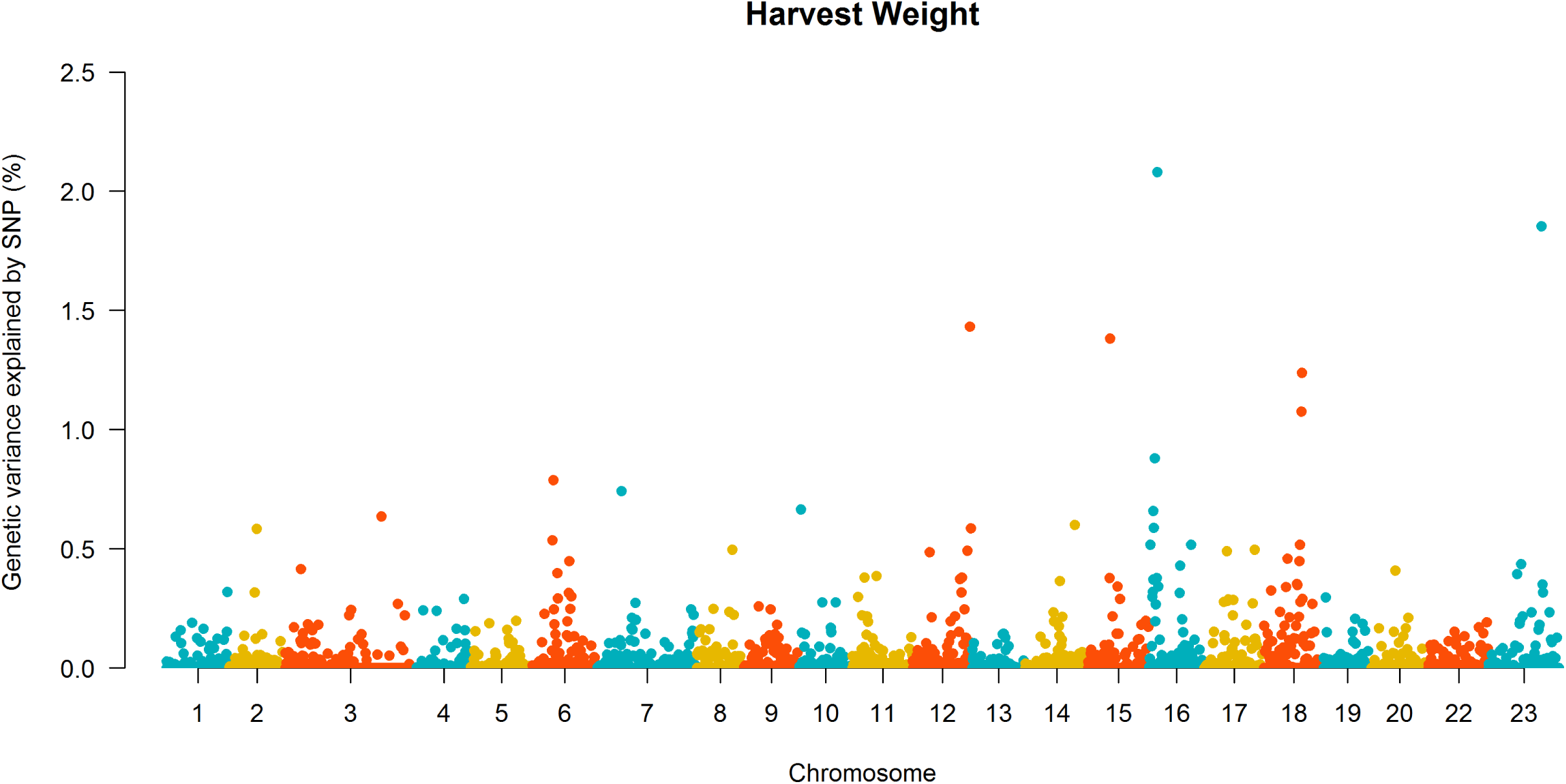
Manhattan plot of genetic variance explained by 20-SNP windows for harvest weight in the 2^nd^ iteration of wssGBLUP.

The full list of genes located within the top five 20-SNPs windows associated with FW and HW is shown in Supplemental Table 1. Some candidate genes found within the top five most important windows have been suggested to be involved with growth-related traits in previous studies. For FY we identified genes U3 small nucleolar RNA-associated protein 6 homolog (*UTP6*), Ras-related protein (*Rab31*) and Follistatin-related protein (*FLRG* or *FSTL3*), located in chromosome 06, 18 and 23, respectively. For HW we identified the genes Natriuretic Peptide Receptor 1 (*NPR1*) and Exostosin Like Glycosyltransferase 3 (*EXTL3*) located in chromosome 22 and 15, respectively.

### Accuracy of genotype imputation

We observed that imputation accuracy decreased with reduced marker density going from LD3K to LD0.5K SNP panels with values ranging from 0.98 to 0.90, respectively (Figure 3). The largest increase in imputation accuracy occurred when increasing SNP density from 0.5K to 1K, with an increase in imputation accuracy of about 6%.

**Figure 3.**
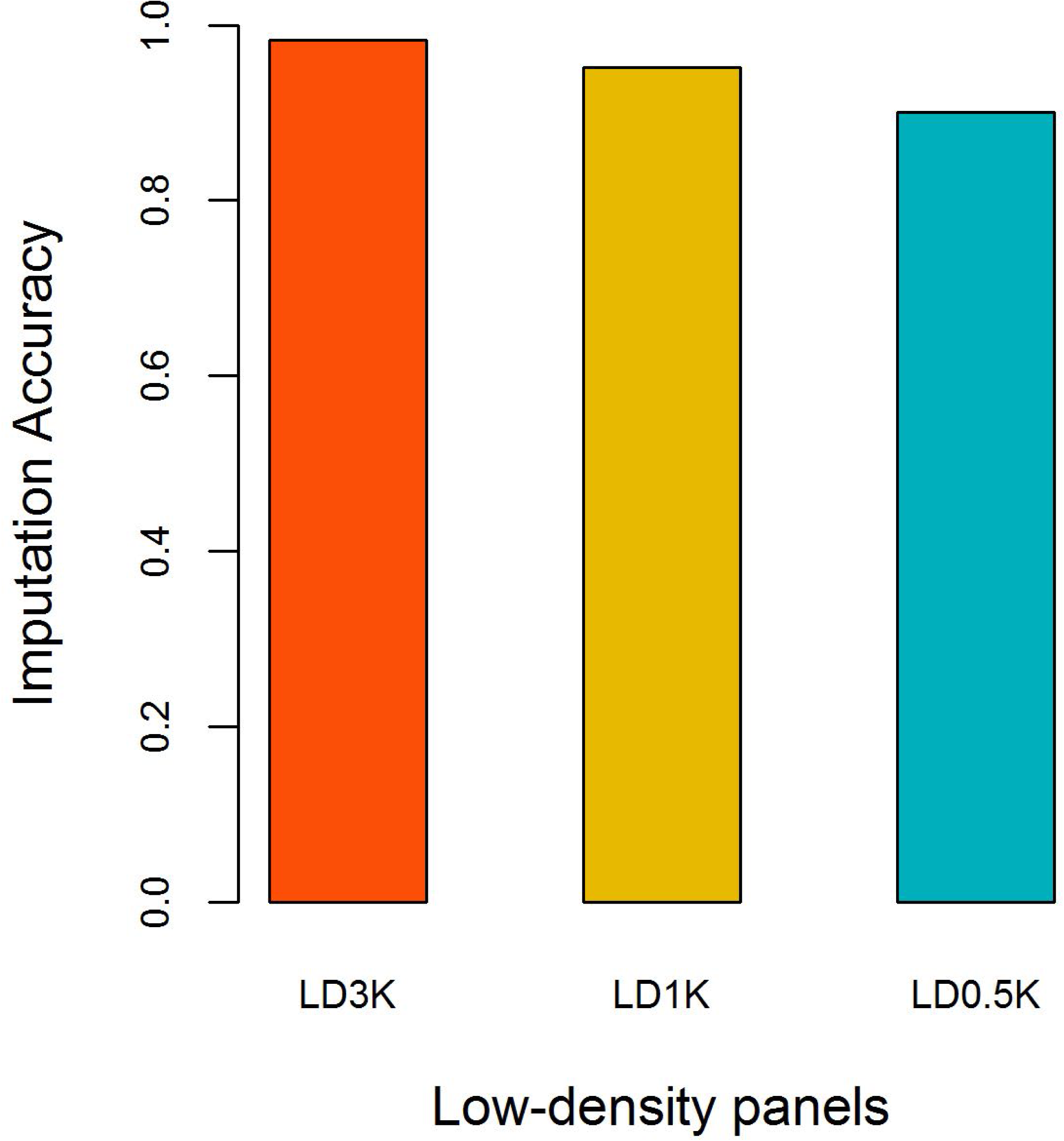
Imputation accuracy from low-density (LD3K, LD1K and LD0.5K) to high-density (HD) panel in Nile tilapia using parents (n = 108) and 20% of offspring (n = 226) genotyped with the HD panel as the reference set and 80% of offspring (n = 904) as the validation set.

Supplemental Figures 1, 2 and 3 show the correlation between observed and imputed genotypes for each SNP in all chromosomes using the LD3K, LD1K and LD0.5 panels, respectively. Imputation accuracy was not consistent across chromosomes, especially for LD0.5K. Inconsistencies may happen because of the physical position of imputed SNP and the location of the SNP on the LD panel. The imputation accuracy decreased greatly at telomeres, and increased considerably with increased SNP density.

### Accuracy of PBLUP and ssGBLUP

Based on the five-fold cross validation, the prediction accuracy for GEBV from genomic methods outperformed the accuracy for EBV from PBLUP. In addition, the accuracy of genomic selection using imputed genotypes from LD to HD SNP panels outperformed both PBLUP and ssGBLUP using true LD genotypes (Table 4).

**Table 4.** Mean accuracy of EBV and GEBV for fillet yield and harvest weight in Nile tilapia using true and imputed genotype SNPs.

| Traits | PBLUP | True genotypes |  |  |  | Imputed genotypes |  |  |
| --- | --- | --- | --- | --- | --- | --- | --- | --- |
|  |  | 32K | 3K | 1K | 0.5K | 3K | 1K | 0.5K |
| **Fillet yield** | 0.539 | 0.621 | 0.612 | 0.574 | 0.560 | 0.621 | 0.620 | 0.585 |
| **Harvest weight** | 0.479 | 0.601 | 0.539 | 0.537 | 0.500 | 0.607 | 0.600 | 0.586 |

The relative increase in accuracy of predicted GEBV compared with EBV from PBLUP varied moderately between the LD panels and traits (Figure 4). Thus, the relative increase in accuracy for FY when comparing ssGBLUP to PBLUP ranged from 4 to 15% for true 0.5 and 32K genotypes, respectively and from 8 to 15% for imputed genotypes using the 0.5 and 3K LD SNP panels respectively. For HW the relative increase in accuracy when comparing ssGBLUP to PBLUP ranged from 4 to 25% for true 0.5 and 32K genotypes, respectively and from 22 to 27% for imputed genotypes using the 0.5 and 3K LD SNP panels respectively. In general, the relative increase in accuracy of predicted GEBV from all true LD SNP panels and imputed genotypes were always better than EBV from PBLUP even at the lowest marker density of 0.5 K for all traits. The relative increase in accuracies when comparing ssGBLUP to PBLUP were almost always higher for HW than for FY, except for the use of true 3K genotypes for prediction of FY (14%) which was slightly higher than for HW (13%) using the same 3K genotypes.

**Figure 4.**
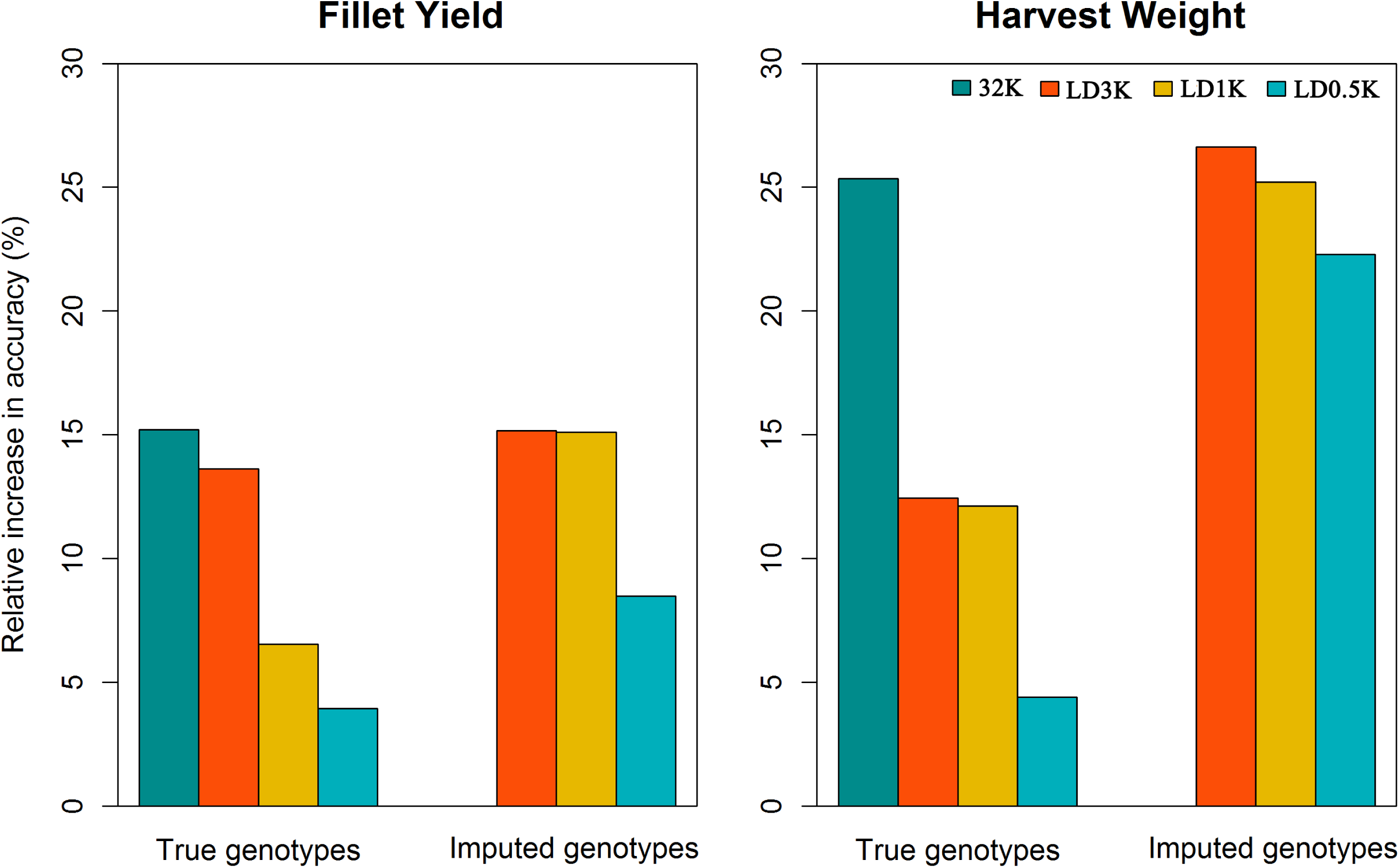
Relative increase in accuracy of different genomic selection methods for fillet yield, harvest weight and waste weight compared to PBLUP in Nile tilapia using true and imputed genotypes.

The genomic prediction accuracy using imputed genotypes, was identical or very similar between the LD panels compared to the 32K SNP genotypes, especially for FY (Figure 4). When comparing the use of true and imputed genotypes for genomic selection it was evident that genotype imputation resulted in a higher increase in relative accuracy independently of trait and LD SNP panel. As expected, the lowest genomic prediction accuracy when using imputed genotypes was always observed for the SNP panel with lowest imputation accuracy (LD0.5K), which resulted in an accuracy slightly lower than the 32K SNP panel, but higher than PBLUP.

## DISCUSSION

### Heritability

We found moderate heritability values for FY and HW which agrees with previous estimates calculated using pedigree-based methods (Gjerde et al., 2012; Nguyen et al., 2010; and Rutten et al., 2005). HW and FY heritability values in tilapia, estimated using genomic information, are reported for the first time in this study. We found slight higher estimates of heritability when using genomic information compared to PBLUP, especially for HW, which is in accordance with what has been reported in other fish species for fillet yield and growth traits (Tsai *et al.* 2015; Gonzalez-Pena *et al.* 2016a).

### Genome-wide association analysis

In the present study, we found no evidence of major quantitative trait loci for both fillet yield and harvest weight in Nile tilapia. The small effect of these loci reinforces evidence of the polygenic nature of these traits. Previous studies support our results and have shown the polygenic nature of fillet yield and growth-related traits in different aquaculture species, with no evidence of major effect genes or genomic regions assessed by GWAS (Gutierrez *et al.* 2015; Tsai *et al.* 2015; Gonzalez-Pena *et al.* 2016a; Yoshida *et al.* 2017; Garcia *et al.* 2018; Reis Neto *et al.* 2019). Furthermore, within the five 20-SNP windows that explained the higher proportion of genetic variance, we found several genes that could potentially be involved in growth and fillet yield.

Although, it is out of the scope of the present study to discuss in detail the putative genes involved in FY and HW, we found it worthy to mention some of the most biologically relevant candidates that may be worthy of functional validation. For instance, the *UTP6* gene is suggested to enhance cellular growth through an increase in the number of ribosomes in Chinese hamster ovary cells (Courtes *et al.* 2013). For both FY and HW, between position 24,563,797 and 24,886,884 bp, we identified the *RAB31* gene, which has a role in trafficking the epidermal growth factor receptor (*EGFR*) gene (Chua and Tang 2014), an important receptor of tyrosine kinase in animals that functions in development, growth and tissue regeneration (Wang *et al.* 2018). In addition, *FSTL3*, present in one of the top 5 SNP windows explaining a high proportion of the genetic variance for FY, is a member of follistatin family, which has been suggested to be an inhibitory binding protein of myostatin activity (Hill *et al.* 2002, 2003). Rebhan and Funkenstein (2008) reported experimental evidence that myostatin activity can be inhibited by follistatin. Chu et al. (2016) observed an increase in the number of muscle fibers, satellite cell activation and decreased expression of myostatin in animals treated with *FSTL3*; suggesting that the gene might be involved to muscle development in the Chinese Perch (*Siniperca chuatsi*).

In plants *NPR1* is an essential regulator of systemic acquired resistance, conferred immunity to broad-spectrum of pathogens (Cao *et al.* 1997; Mou *et al.* 2003). However, Vanacker *et al.* (2001) reported a novel function for NPR1, which is associated with growth control, cell division and suppressing endoreduplication during leaf development in *Arabidopsis*. In humans, a mutation affected the *EXTL3* gene causing skeletal dysplasia, immune deficiency and development delay. In zebrafish abnormalities of cartilage development and defective formation in fin and branchial arch were reported (Norton *et al.* 2005). Other genes located in a 20-SNP window flanking the top five windows are presented in Supplemental Table 1.

### Accuracy of genotype imputation

The imputation accuracy, on average, was above 90%, independent of the LD SNP panel used but decreased from LD3K to LD0.5K, which is in accordance with the same pattern seen by Habier et al. (2009), Hickey et al. (2012) and Yoshida *et al.* (2018b). The imputation errors could be higher in LD panels because they could be less efficient in capturing the linkage and linkage disequilibrium between the markers. Like previous studies we found accuracies of genotype imputation to be very similar using panels of 3K SNPs or denser (Druet *et al.* 2010; Zhang and Druet 2010; Duarte *et al.* 2013; Carvalheiro *et al.* 2014; Cleveland and Hickey 2014; Kijas *et al.* 2016; Tsai *et al.* 2017). However, it is likely that 3K or denser SNP panels will be considerably more expensive than 0.5K or 1K SNP panels, thus cost-effectiveness must be carefully evaluated in further studies.

Some imputation studies tested the size of the reference population (Zhang and Druet 2010; Cleveland and Hickey 2014; Tsai *et al.* 2017; Yoshida *et al.* 2018b), and have shown that the number of animals used in this study should be sufficient to not influence the imputation accuracy. Therefore, we did not include the effect of different genotyping strategies in the final results; however, in preliminary tests we found imputation accuracies lower than 90% using a small proportion of offspring (less than 10%) in the reference set when imputing genotypes from the 0.5K SNP panel (results not show). This is probably because of the small number of animals per family in the reference population, which can influence imputation accuracy (Hickey *et al.* 2012). In this case we had approximately 18 sibs genotyped/family and inclusion of 20% of offspring in addition to the parents genotyped with the 32K SNP panel as the reference set, to achieve similar accuracy values to those reported by Yoshida *et al.* (2018b) for *Salmo salar* where 31 sibs/family and just 10% of offspring were needed to surpass an imputation accuracy of 90%. The influence of the small number of animals per family in the reference set decreased when the density of the LD SNP panels increased (Carvalheiro *et al.* 2014).

Supplemental Figures 1, 2 and 3 indicate regions of the genome containing markers with high imputation errors, especially at the beginning and end of the chromosomes. This is could be an effect of recombination rates, that are known to be higher around the telomeres (Chowdhury *et al.* 2009; Tortereau *et al.* 2012). The physical location of the SNP is another factor that has been shown to be affect the imputation accuracy and to reduce the errors in these regions, some previous studies suggested increasing the coverage of SNP chromosomal extremes (Badke *et al.* 2012; Boichard *et al.* 2012; Dassonneville *et al.* 2012). In addition, high imputation errors far from chromosome extremes can be the result of erratic patterns of linkage disequilibrium, which suggests potential issues related to physical maps and reference genome assembly (Druet *et al.* 2010; Carvalheiro *et al.* 2014; Yoshida *et al.* 2018b).

Another important factor that may affect the imputation accuracy is the linkage disequilibrium between markers which is exploited to infer the missing genotypes (Hickey *et al.* 2012; Carvalheiro *et al.* 2014). A previous study, showed a more rapid decrease of linkage disequilibrium with inter-marker distance for this Nile tilapia population (Yoshida *et al.* 2019) when compared to other populations of different aquaculture species (Barria et al., 2018b; Gutierrez et al., 2015; Kijas et al., 2016; Vallejo et al., 2018). Nevertheless, our imputation accuracies are close to the imputation values reported in the literature for salmonids (Kijas *et al.* 2016; Tsai *et al.* 2017; Yoshida *et al.* 2018b) and terrestrial species (Badke *et al.* 2012; Hayes *et al.* 2012; Duarte *et al.* 2013; Hozé *et al.* 2013; Carvalheiro *et al.* 2014), suggesting that the family-based imputation approach is less sensitive to linkage disequilibrium patterns by efficiently exploiting information of highly related animals.

### Accuracy of genomic prediction

Our results showed that the use of genomic information for estimating breeding values achieved higher accuracies compared to using only pedigree information for FY and HW, independent of the LD SNP panel used, with or without imputation of genotypes (Figure 4). The relative increase in GEBV accuracies compared to PBLUP has been previously reported for growth (Garcia et al., 2018; Tsai et al., 2015) and for different disease resistance traits in farmed aquaculture species (Tsai *et al.* 2016; Vallejo *et al.* 2017; Bangera *et al.* 2017; Correa *et al.* 2017; Barria *et al.* 2018a; Yoshida *et al.* 2018a, 2018c).

The accuracy of GEBV depends on factors such as the number of genotyped and phenotyped individuals in the training population, the heritability and the number of loci affecting the trait (Daetwyler *et al.* 2008; Goddard 2009). Furthermore, the accuracy of genomic prediction is highly dependent on the genotype density used, which means that increasing marker densities tends to generate higher GEBV accuracies (Tsai *et al.* 2016; Bangera *et al.* 2017; Correa *et al.* 2017; Yoshida *et al.* 2018a). In addition, the use of different methods to estimate the GEBV can directly affect the accuracy of genomic prediction. In general, the genomic methods differ in distributional assumptions of marker effects and the calculation of the genetic relationship matrix. Here, we used the ssGBLUP method (Misztal *et al.* 2009), which assumed a normal distribution of marker effects and has some of practical advantages, given that it uses information from genotyped and nongenotyped animals (Lourenco *et al.* 2014), and it has also been demonstrated to provide higher accuracy than the PBLUP method and other genomic methods (Chen *et al.* 2011; Christensen *et al.* 2012; Vallejo *et al.* 2017; Yoshida *et al.* 2018a).

To test the impact of genotype imputation errors in genomic predictions we estimated the accuracy of genomic predictions for FY and HW using imputed genotypic data and compared the data to true 32K and LD SNP genotypes (LD3K, LD1K and LD0.5K). Our results indicate that genomic prediction accuracies using imputed genotypes were always higher than those obtained using true LD genotypes and equal or slightly lower than using true 32K genotypes (Figure 4). For HW the accuracy of GEBV was substantially higher using the imputed LD0.5K than true LD0.5K, LD1K and even LD3K, whereas for FY the accuracy using imputed LD0.5K did not surpass the true LD3K panel. Our results are in accordance with previous studies in aquaculture (Dufflocq et al., 2019; Tsai et al., 2017; Yoshida et al., 2018b) and livestock species (Berry and Kearney 2011; Erbe *et al.* 2012); indicating that the use of genotype imputation can decrease the cost of genotyping by means of using less expensive LD SNP panels without compromising prediction accuracies. In addition, the influence of imputation error on genomic prediction accuracy depends on the genetic architecture underlying the studied traits. For traits that are influenced by few QTLs with large effect, accuracy of genomic prediction could be more sensitive to imputation errors than polygenic traits, such FY and HW (Chen *et al.* 2014).

### Implications

The cost associated with genotyping represents a key limiting factor for the practical implementation of genomic selection in fresh water fish species. One of the main objectives of this study was to test genotyping imputation as an alternative to reduce the costs of genotyping for genomic selection in Nile tilapia breeding programs. Dufflocq et al. (2019) and Tsai et al. (2017) suggested that for aquaculture species the use of imputation strategies can reduce the cost of genotyping by at least 60% compared to genotyping all animals with a high-density panel. The genotype cost depends on the density of the SNP panel, genotyping technology and number of samples. In general the price ranges from USD $5 to USD $75 for low-(0.5K) and high-density panels (50K), respectively. The strategies used in present study for genotyping both parents and a proportion of offspring resulted in low imputations errors; suggesting that genotype imputation represents a promising strategy for the practical application of genomic selection in tilapia.

## CONCLUSIONS

The GWAS indicated a polygenic architecture for fillet yield and harvest weight, with some markers explaining a small proportion of genetic variance; indicating that the implementation of marker assisted selection could not be successfully applied for these traits in the present Nile tilapia population. In contrast, the use of genomic selection could increase the response to selection and improve genetic progress. The use of genotype imputation can reduce genotyping costs and allow the implementation of genomic selection in Nile tilapia breeding programs.

## ACKNOWLEDGEMENTS

This work has been funded by Corfo (project number 14EIAT-28667). The authors are grateful to Acuacorporación Internacional for providing the Nile tilapia dataset and fin samples.

## AUTHORS’ CONTRIBUTIONS

GMY performed the analysis and wrote the initial version of the manuscript. KC and JPL contributed with study design. JS and DS contributed to family formation and data collection. JMY conceived and designed the study; contributed to the analysis, discussion and writing. All authors have reviewed and approve the manuscript.

---

**Supplemental Table 1.**
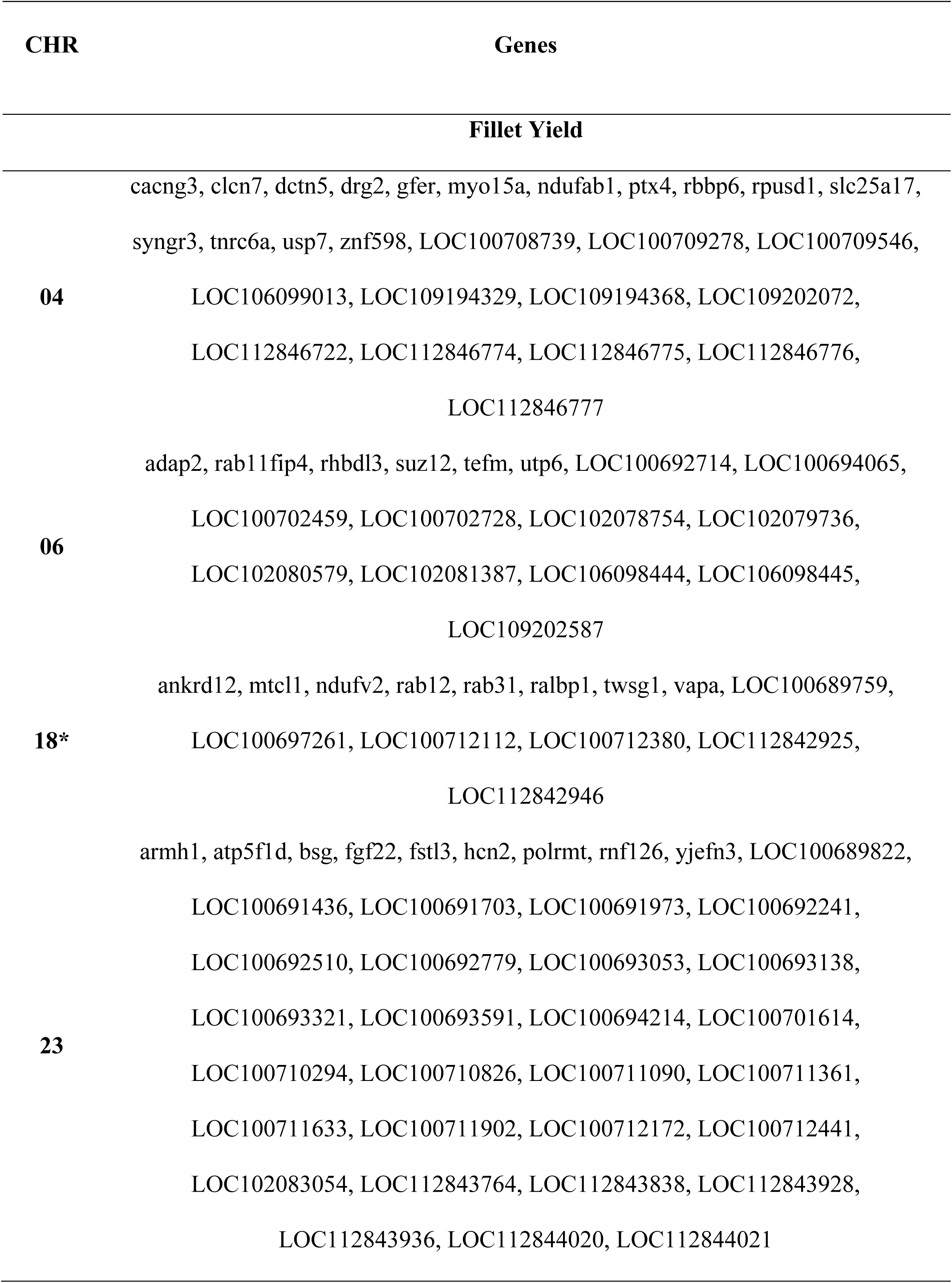
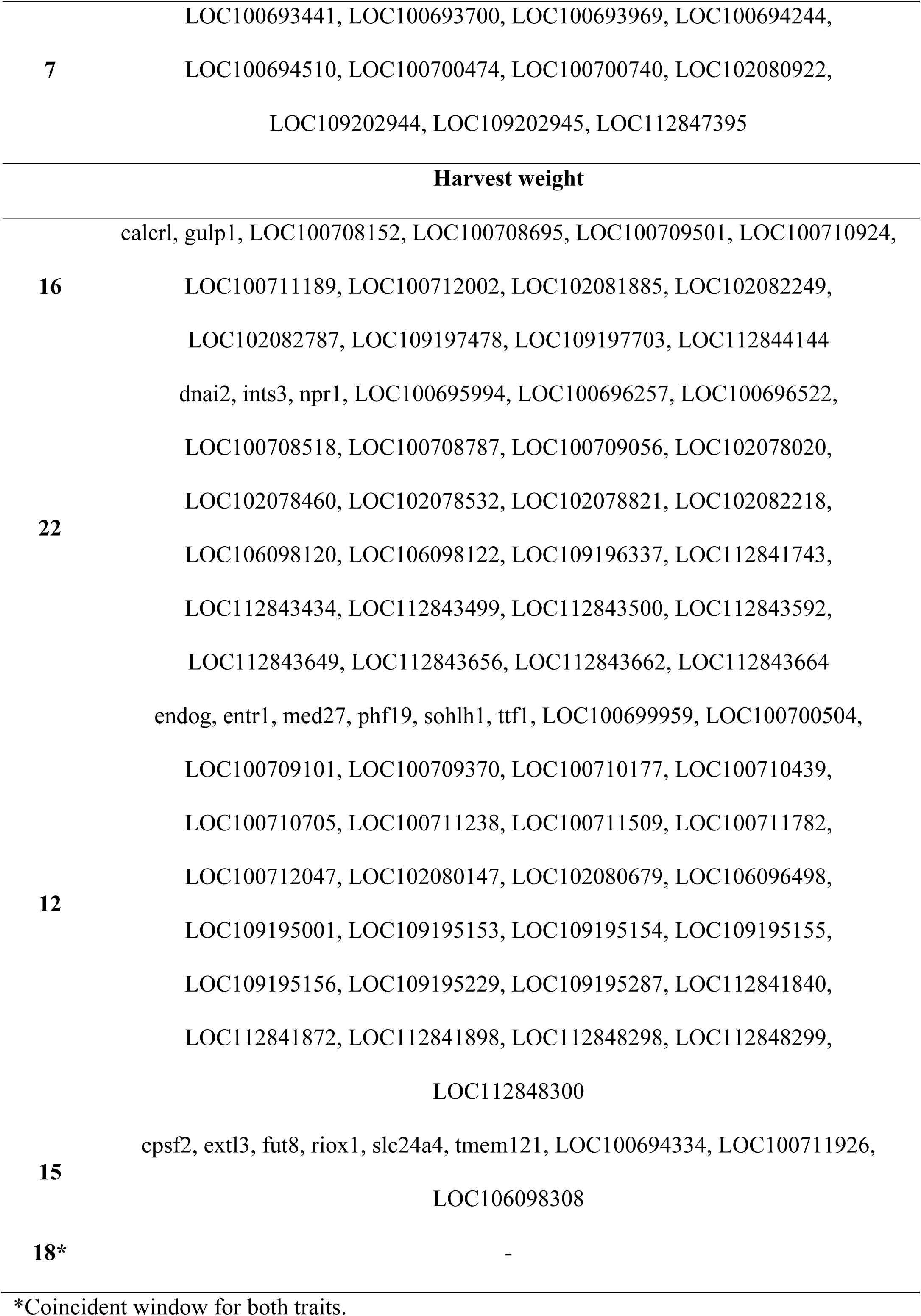
Full list of genes located within the top five 20-SNP windows associated with fillet yield and harvest weight in Nile tilapia.

**Supplemental Figure 1.**
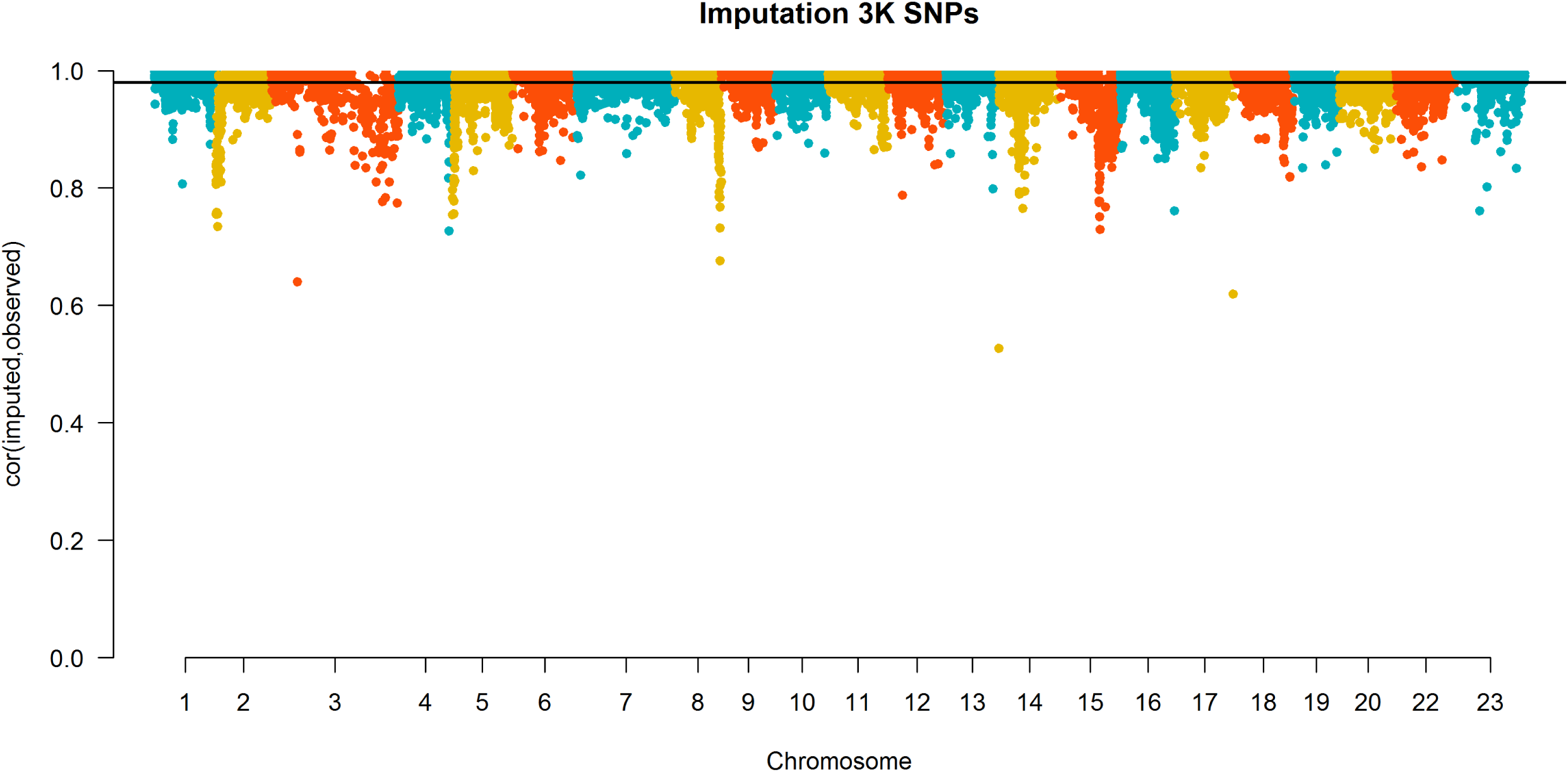
Correlations between observed and imputed genotypes for each SNP for imputation from low-density (LD3K) to high-density (HD) panel in Nile tilapia using parents (n = 108) and 20% of offspring (n = 226) genotyped with the HD panel as the reference set and 80% of offspring (n = 904) as the validation set. The black line indicates the mean imputation accuracy.

**Supplemental Figure 2.**
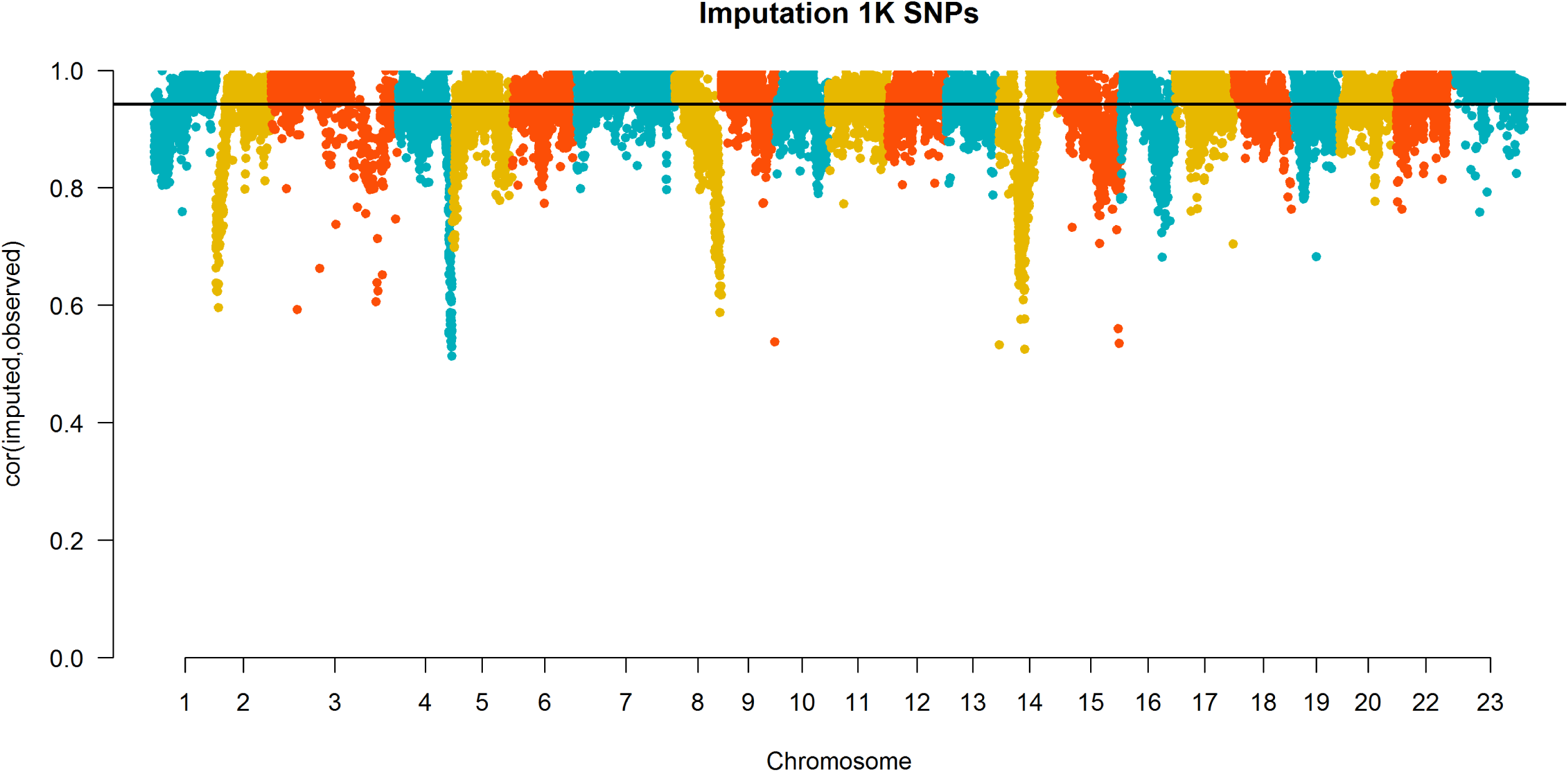
Correlations between observed and imputed genotypes for each SNP for imputation from low-density (LD1K) to high-density (HD) in Nile tilapia using parents (n = 108) and 20% of the offspring (n = 226) genotyped with the HD panel as the reference set and 80% of the offspring (n = 904) as the validation set. The black line indicates the mean imputation accuracy.

**Supplemental Figure 3.**
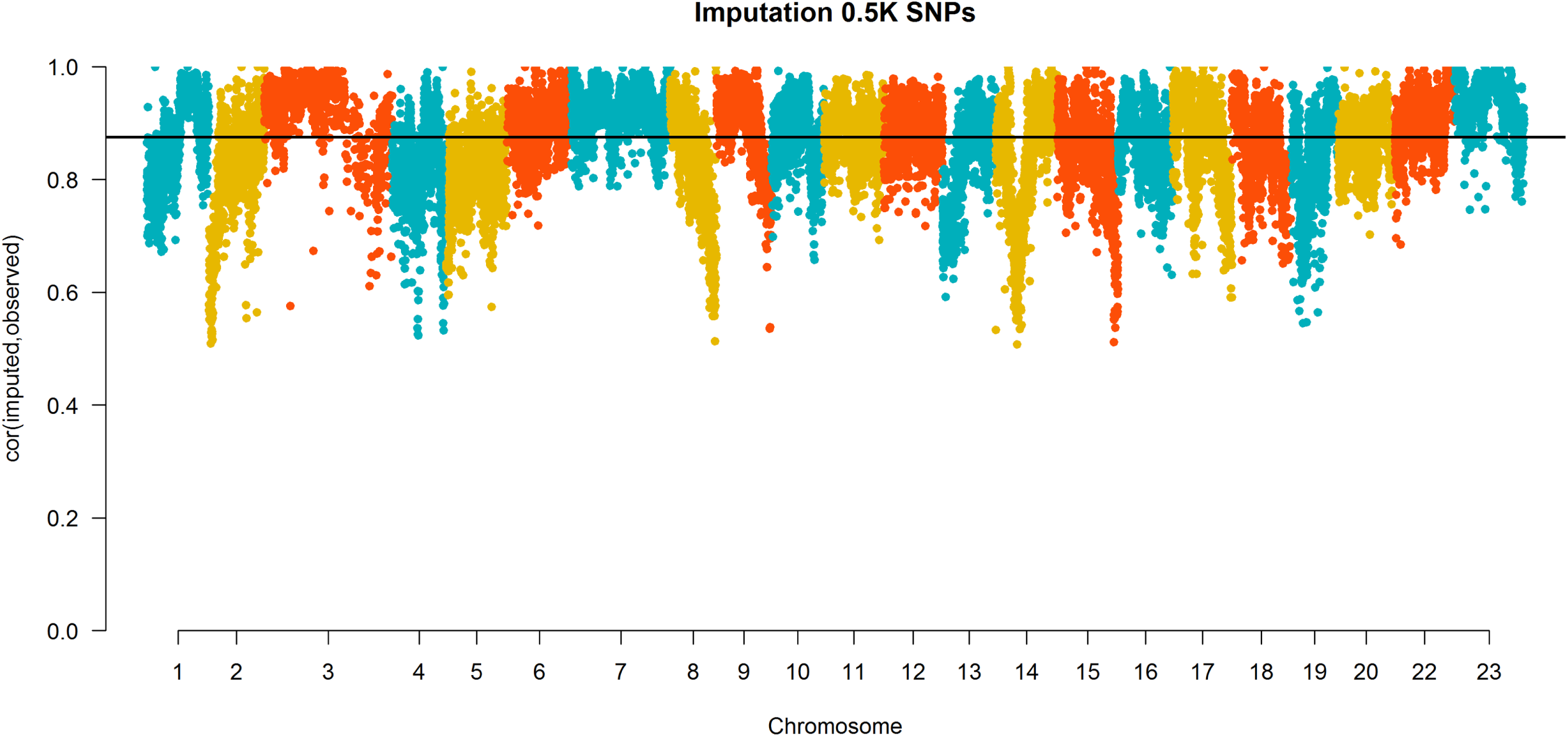
Correlations between observed and imputed genotypes for each SNP for imputation from low-density (LD0.5K) to high-density (HD) in Nile tilapia using parents (n = 108) and 20% of offspring (n = 226) genotyped with the HD panel as the reference set and 80% of the offspring (n = 904) as the validation set. The black line indicates the mean imputation accuracy.

